# Production and purification of human Hsp90β in *Escherichia coli*

**DOI:** 10.1101/121996

**Authors:** Martina Radli, Dmitry B. Veprintsev, Stefan G. D. Rüdiger

## Abstract

The molecular chaperone Hsp90 is an essential member of the cellular proteostasis system. It plays an important role in the stabilisation and activation of a large number of client proteins and is involved in fatal disease processes e.g. Alzheimer disease, cancer and cystic fibrosis. This makes Hsp90 a crucial protein to study. Mechanistic studies require large amounts of protein but the production and purification of recombinant human Hsp90 in *E. coli* is challenging and laborious. Here we identified conditions that influence Hsp90 production and optimised a fast and efficient purification protocol. We found that the nutrient value of the culturing medium and the length of induction had significant effect on Hsp90 production in *Escherichia coli*. Our fast, single-day purification protocol resulted in a stable, well-folded and pure sample that was resistant to degradation in a reproducible manner. We anticipate that our results provide a useful tool to produce higher amount of pure, well-folded and stable recombinant human Hsp90β in *Escherichia coli* in an efficient way.

## Introduction

The cellular proteostasis system evolved to maintain cellular health and stability and to protect cells from continuously occurring stress by a tight control of protein production, quality control, folding, trafficking, aggregation and degradation (1,2). Chaperones are crucial elements of the proteostasis system, they prevent protein misfolding and aggregation by various mechanisms and thereby contribute to cellular integrity (3).

The molecular chaperone Hsp90 is one of the most important element of the proteostasis system (4-7). It is involved in client protein folding, maturation, stabilisation, activation and assembly of large protein complexes (8,9). Typically, Hsp90 assists at late folding processes (10,11).

Hsp90 interacts with up to 10 % of the cellular proteome. Its main clients include transcription factors, kinases and hormone receptors (Didier Picard. Table of Hsp90 interactors. Available from: https://www.picard.ch/downloads/Hsp90interactors.pdf). Moreover, it has unconventional partners such as the disordered α-synuclein and Tau (5,10). Hsp90 plays a crucial role in the progression of several diseases (such as cancer, cystic fibrosis and Alzheimer’s disease) (12-14).

The Hsp90 chaperone machinery is extensively studied to understand its complex working mechanism and involvement with multiple fatal diseases. Typically, *in vitro* biochemical and biophysical experiments require high amount of recombinant protein with excellent sample purity and stability. Significant overproduction of the recombinant human Hsp90β is challenging since it is a large, multi-domain protein. Moreover, Hsp90 is particularly sensitive to the proteolytic cleavage of the flexible, unfolded linkers between its domains, therefore a time consuming purification protocol may compromise sample quality.

We systematically tested several parameters to find the best conditions for the overproduction of recombinant human Hsp90β. Furthermore, we developed a fast and efficient purification protocol that results in pure and stable sample with reproducible quality. We found that the nutrient value of the medium and the length of induction time had significant effect on Hsp90 overproduction, whereas the concentration of induction agent (isopropyl β-D-1-thiogalactopyranoside (IPTG)), temperature and optical density (OD_600_) at induction (within the tested intervals) did not influence the process. The protein samples we purified with the optimised protocol were resistant to proteolysis upon incubation at physiological temperature up to one day. We showed that the folding, sedimentation and molecular weight of our Hsp90 sample corresponded to earlier results.

## Materials and Methods

### Wild type full length Hsp90 construct

pet23a+ expression vector (Novagen) was used for His_6_-tagged wild type human Hsp90 production in Rosetta 2 *E. coli* strain (Novagen). The sequence of wild type Hsp90:

HHHHHHMPEEVHHGEEEVETFAFQAEIAQLMSLIINTFYSNKEIFLRELISNASDALDKIRYESLTDPSKLDSGKELKIDIIPNPQERTLTLVDTGIGMTKADLINNLGTIAKSGTKAFMEALQAGADISMIGQFGVGFYSAYLVAEKVVVITKHNDDEQYAWESSAGGSFTVRADHGEPIGRGTKVILHLKEDQTEYLEERRVKEVVKKHSQFIGYPITLYLEKEREKEISDDEAEEEKGEKEEEDKDDEEKPKIEDVGSDEEDDSGKDKKKKTKKIKEKYIDQEELNKTKPIWTRNPDDITQEEYGEFYKSLTNDWEDHLAVKHFSVEGQLEFRALLFIPRRAPFDLFENKKKKNNIKLYVRRVFIMDSCDELIPEYLNFIRGVVDSEDLPLNISREMLQQSKILKVIRKNIVKKCLELFSELAEDKENYKKFYEAFSKNLKLGIHEDSTNRRRLSELLRYHTSQSGDEMTSLSEYVSRMKETQKSIYYITGESKEQVANSAFVERVRKRGFEVVYMTEPIDEYCVQQLKEFDGKSLVSVTKEGLELPEDEEEKKKMEESKAKFENLCKLMKEILDKKVEKVTISNRLVSSPCCIVTSTYGWTANMERIMKAQALRDNSTMGYMMAKKHLEINPDHPIVETLRQKAEADKNDKAVKDLVVLLFETALLSSGFSLEDPQTHSNRIYRMIKLGLGIDEDEVAAEEPNAAVPDEIPPLEGDEDASRMEEVD

### Protein overproduction test - cell culturing

Rosetta 2 cells containing pet23a+ vector with wild type full length Hsp90 were inoculated into 100 ml 2x yeast tryptone extract (2x YT; 12.8 g Bacto Tryptone (Merck), 8 g Bacto Yeast Extract (Merck), 4 g NaCl (Merck) in 800 ml demi water (Millipore)) medium supplemented with 34 mg/l final concentration of chloramphenicol (Sigma-Aldrich) and 100 mg/l final concentration of ampicillin (Sigma-Aldrich). The cells were grown over night at 37°C, shaking with 220 rpm.

Next morning 100 (for low OD_600_) or 200 (for high OD_600_) μl culture was inoculated into 4 ml lysogeny broth medium (LB; 8 g Bacto Tryptone, 4 g Bacto Yeast Extract, 4 g NaCl, 20 g tymine (Sigma-Aldrich) in 800 ml demi water), 2x YT or terrific broth media (TB; 12 g Bacto Tryptone, 24 g Bacto Yeast Extract, 4 ml glycerol in 800 ml demi water)) supplemented with 34 mg/l final concentration of chloramphenicol and 100 mg/l final concentration of ampicillin. The cultures were grown at 37°C, shaking with 180 rpm. The OD_600_ was monitored every 1.5 hours by using a 1:10 dilution of the culture in an Ultrospec 3000 pro UV/Visible Spectrophotometer (GE Healthcare). The cultures were induced at low (0.6-0.9) or high (1-1.3) OD_600_ with 0.1/0.25/0.5 mM IPTG (Thermo Fisher Scientific). The cultures were incubated at 16°C or 18°C, the SDS-PAGE samples were taken before induction and 1/3/5 day(s) after induction.

### SDS-PAGE

For the SDS-PAGE samples 0.5 ml of cell culture was taken and centrifuged for 1 minute by 13,300 rpm. The supernatant was discarded and the pellet was resuspended in 200 μl of 1x sample buffer (0.625 M Tris (Sigma-Aldrich), 12.5 % glycerol (CARL ROTH), 1 % SDS (Bio-Rad), 0.005 % Bromophenol Blue (Bio-Rad), 5 mM freshly added β-mercaptoethanol (Sigma-Aldrich) 2x diluted with demi water (Millipore)) and homogenised by using a syringe of a very tiny diameter (BD Micro-Fine). This last step was essential for loading the samples on the gel.

15 % SDS gels were prepared (Separation buffer: 0.38 M Tris pH 8.8, 15 % acrylamide (National Diagnostics), 0.1 % SDS, 0.1 % APS (Sigma-Aldrich), 0.04 % TEMED (Sigma-Aldrich), Stacking buffer: 0.125 M Tris pH 6.8, 4 % acrylamide, 0.1 % SDS, 0.075 % APS, 0.1 % TEMED) and ran in 1x Laemmli buffer (0.025 M Tris base, 0.152 M glycine (SERVA Electrophoresis GmbH), 0.1 % SDS, diluted from 10x stock). The gels stained with Coomassie staining solution (0.2 % Coomassie Brilliant Blue (SERVA Electrophoresis GmbH), 45 % methanol (Interchema Antonides-Interchema), 10 % acetic acid (Biosolve) and 55 % demi water and destained using destaining solution (30 % methanol, 10 % acetic acid and 60 % demi water).

The gels were scanned in a Epson Perfection V700 Photo scanner. Quantification of the lane profiles was done by using ImageJ. The lane profiles of the induced samples were normalised to the lane profile of the uninduced sample of the same gel using the intrinsic *E. coli* protein called EF-Tu. Subsequently, the lane profiles of induced samples were aligned to the uninduced sample using the first two bands in the beginning of the lanes. Finally, the uninduced lane profile was deducted from the induced lane profiles and the sum of each Hsp90 peak area was divided by the sum of the normalised EF-Tu peak area of the same lane to be able to compare the gels with each other.

### Overproduction and purification of wild type full length Hsp90

Rosetta 2 cells containing pet23a+ vector with wild type full length Hsp90 were inoculated into 200 ml 2x YT medium supplemented with 34 mg/l final concentration of chloramphenicol and 100 mg/l final concentration of ampicillin. The cells were grown over night at 37°C, with shaking at 220 rpm. Next morning 200 ml culture was inoculated into 6x800 ml 2x YT supplemented with 34 mg/l final concentration of chloramphenicol and 100 mg/l final concentration of ampicillin. The cultures were grown at 37°C with shaking at 180 rpm and induced with 0.5 mM IPTG at OD_600_ = 1. After induction the cells were incubated at 18°C for 5 days with shaking at 180 rpm.

The cells were harvested in an Avanti J-26 XP centrifuge (Beckman Coulter) using the JLA-8.1 rotor at 4°C at 4500 rpm for 30 minutes. The supernatant was discarded and the pellet was resuspended in ice cold resuspension-buffer (50 mM Na-phosphate pH 7.2 (Sigma-Aldrich), 150 mM NaCl, 150 mM KCl (CARL ROTH)) and centrifuged in an MSE Harrier 18/80 Refrigerated Benchtop Centrifuge at 4°C at 5000 rpm for 30 minutes. The supernatant was discarded and the pellet was stored at -20°C until further usage.

The pellet was resuspended in ice cold lysis buffer (12.5 mM Na-phosphate pH 7.2. 75 mM NaCl, 5 mM β-mercaptoethanol, EDTA-free protease inhibitor (1 tablet/50 ml) (Roche)). The cells were disrupted by an EmulsiFlex-C5 (Avestin) cell disruptor. The lysate was centrifuged in Avanti J-26 XP centrifuge using JA-25.5 rotor at 21,000 rpm for 45 minutes at 4°C. The lysate was filtered by 22 μm polypropylene filter (VWR) to remove the cell debris and insoluble aggregates. The purification was done using an AKTA Purifier (GE Healthcare).

Wild type full length Hsp90 was first purified on an IMAC POROS 20MC (Thermo Fischer Scientific) affinity purification column (solutions connected to pump A1 and A2: 50 mM Na-phosphate buffer pH 8.0 with 300 mM NaCl, B1: demi water with 10 mM β-mercaptoethanol, B2: 1 M imidazole (Sigma-Aldrich)). The eluted sample was diluted 4-fold with dilution buffer (25 mM Na-phosphate buffer pH 7.2, 5 mM DTT (Sigma-Aldrich), complete protease inhibitor (1 tablet/50 ml) (Roche)). Next the sample was loaded on a POROS 20HQ anion exchange column (Thermo Fischer Scientific) (solutions connected to pump A1 and A2: 50 mM Na-phosphate pH 7.2, B1: demi water with 10 mM DTT, B2: 2 M KCl). The eluted sample was diluted 10-fold with dilution buffer (25 mM Na-phosphate buffer pH 7.2, 5 mM DTT, complete protease inhibitor (1 tablet/100 ml)). Finally, the sample was loaded on a HiTrap heparin affinity chromatography column (GE Healthcare) (solutions connected to pump A1 and A2: 25 mM Na-phosphate pH 7.2, B1: demi water with 10 mM DTT, B2: 2 M KCl). The eluate was concentrated and buffer exchanged to Hsp90 storage buffer (25 mM Na-phosphate pH 7.2, 150 mM NaCl, 150 mM KCl), 5 mM DTT, complete protease inhibitor (1 tablet/100 ml)) using a Vivaspin 20 column (50 kDa MWCO) (GE Healthcare) at 4°C at 5000 rpm for 15-15 minutes until above 100 μM protein concentration. The protein concentration was determined with ND-1000 program on an ND-1000 Spectrophotometer type NanoDrop using 57760 M^−1^cm^−1^ extinction coefficient. The sample was aliquoted, frozen in liquid N_2_ and stored at -80°C. Throughout the purification procedure samples were taken from the steps of the purification that were run on SDS-PAGE to analyse sample purity.

### Protein stability

10 μM concentration Hsp90 was incubated in Hsp90-buffer ((25 mM Na-phosphate pH 7.2, 150 mM NaCl, 150 mM KCl, 5 mM DTT, complete protease inhibitor (1 tablet/100 ml)) for 24 hours at 4°C, room temperature (∼21°C) and 37°C. SDS-PAGE samples were taken and mixed with 2x sample buffer (1.25 M Tris, 25 % glycerol, 2 % SDS, 0.01 % Bromophenol Blue, 5 mM freshly added DTT) at 0/3/18/24 hours. The samples were analysed on SDS-PAGE.

### Silver staining

The Coomassie-stained and destained gels were fixed for 30 minutes in fixation solution (30 % ethanol (Interchema Antonides-Interchema), 10 % acetic acid, 60 % demi water). The gel was washed 3-times for 20 minutes in 50 % ethanol, then pre-treated for 1 minute with 0.02 % Na_2_S_2_O_3_ (Scharlau Chemicals) [100x diluted from stock (stock: 2 g into Na_2_S_2_O_3_ 100 ml water (Milli-Q))] and quickly rinsed 4 times with demi water. The gels were impregnated for 20 minutes with freshly prepared staining solution (2 g/l AgNO_3_ (Merck) and 0.75 ml/l formaldehyde (Calbiochem)). Next they were quickly rinsed 4 times with demi water, then developed till the desired result in developer solution (60 g/l Na_2_CO_3_ (Sigma-Aldrich), 0.5 ml/liter formaldehyde and 0.0004 % Na_2_S_2_O_3_ stock). The reaction was stopped in fixation solution (30 % ethanol, 10 % acetic acid, 60 % demi water) for 10 minutes. The gel was stored in 1 % acetic acid solution and scanned by Epson Perfection V700 Photo scanner.

### CD spectroscopy

The protein was centrifuged on 4°C for 15 minutes at 13,300 rpm in Heraeus Pico 17 centrifuge (Thermo Scientific). The concentration was determined with ND-1000 program on ND-1000 Spectrophotometer type NanoDrop using 57760 M^−1^cm^−1^ extinction coefficient. The protein was diluted to 0.1 g/l concentration with CD-buffer (25 mM Na-phosphate pH 7.2 buffer, 150 mM NaF (Sigma-Aldrich)), loaded into a Teflon-sealed, polarimetrically checked quartz glass cuvette with an optical path length of 1 mm and a volume of 350 μl (Hellma Analytics). Far-UV CD spectrum was measured with the Spectra Manager (Jasco) program on a Jasco J-810 Spectropolarimeter instrument (Jasco). Experimental parameters included a wavelength increment of 1 nm, a scan speed of 20 nm/min, a temperature of 20°C). The data were analysed by MS Excel.

### SEC-MALLS

The protein sample was centrifuged and its concentration was measured as described in the *CD spectroscopy* part. 10 μl of ∼29 g/l sample was run with Shimadzu on a Superdex 200 column with 0.35 ml/min speed in running buffer (25 mM Na-phosphate pH 7.2, 150 mM NaCl, 5 mM freshly added DTT). The 220 nm absorption was detected by SPD-20A UV detector (Shimadzu), light scattering by Wyatt COMET^TM^ light scattering detector (Wyatt Technology) and the refractive index by the RID-10A refractive index detector (Shimadzu). The data were analysed by program Astra 6.

### Analytical Ultracentrifugation (*AUC*)

The protein sample was centrifuged and its concentration was measured as described in the CD *spectroscopy* part. The protein sample was diluted to 7.2 μM concentration with Hsp90-buffer (25 Na-phosphate pH 7.2, 150 mM NaCl, 150 mM KCl, complete protease inhibitor (1 tablet/50 ml), 5 mM freshly added DTT). The sample was centrifuged on 20°C for 16 hours at 42,000 rpm in a Beckman XL-I ultracentrifuge using An60Ti rotor. We used absorbance detection optics for our experiment. The data was analysed by SedFit (Schuck, 2000).

## Results

### Systematic testing of recombinant human Hsp90β *overproduction*

We set out to systematically test the effect of key parameters on recombinant human Hsp90β production in Rosetta 2 cells and to identify conditions resulting in high protein levels.

We outlined our study as follows: We cultured *E. coli* cells containing human Hsp90β open reading frame on a pET23a+ vector and tested media with different nutrient values and different temperature of induction to modulate the metabolism of *E. coli* cells. We induced the cultures with different final concentration of IPTG affecting the level of T7 polymerase. We also varied the length of induction and the OD_600_ value at induction, because in case of certain proteins these are essential parameters for successful overproduction.

At given time points we took samples from the cultures and ran them on SDS-gels (**Figure 1A**). We loaded protein marker in the first, uninduced sample in the second and purified Hsp90 in the last lanes of the SDS-gels. These samples helped to identify the Hsp90 bands that appeared after induction. Lane 3-14 contained the induced samples of the different conditions tested. The most abundant intrinsic *E. coli* band that we later used for quantification (EF-Tu) and Hsp90 bands are marked on the right of the gels (**Figure 1A**) (15).

**Figure 1:**
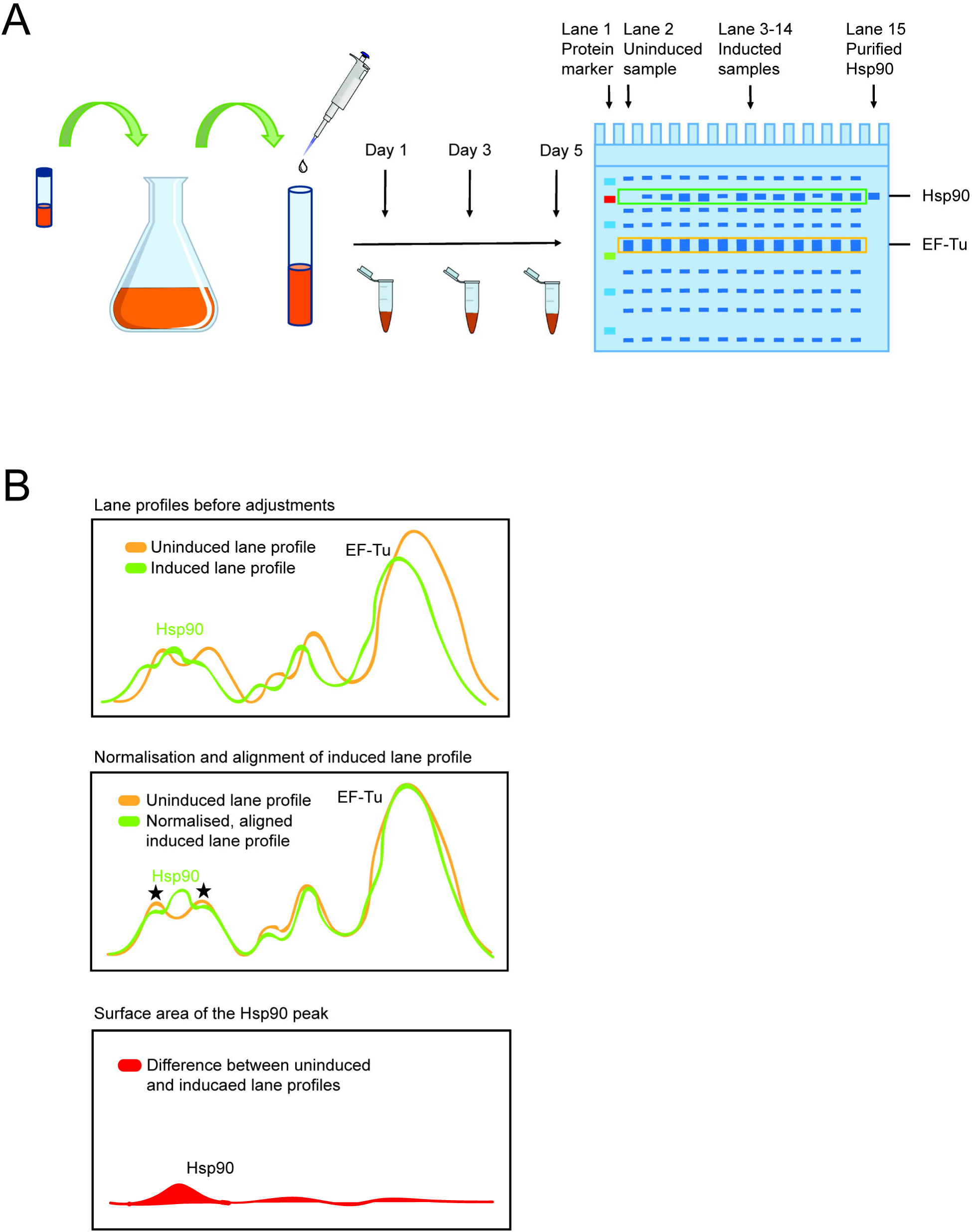
Experimental set up and quantification scheme for Hsp90 overproduction. (A) Schematic overview of the protein production experiment. E. coli cells containing human Hsp90β open reading frame were inoculated from glycerol stock and cultured overnight at 37°C. The culture was divided into small tubes and after induction the tubes were incubated in different conditions. Gel samples were taken at day 1, 3 and day 5 and were run on SDS-PAGE. (B) Schematic representation of the quantification of Hsp90 overproduction. The intensity profiles of uninduced (orange) and induced lanes (green) were determined and the induced lane profile were normalised to the uninduced lane profile using the EF-Tu peak of the uninduced lane as reference (100 %) (top panel). Subsequently, the induced lane profiles were aligned to the uninduced lane profile using the first two left peaks (stars) of the uninduced lane profile as reference points (middle panel). The uninduced lane profile was deducted from the normalised and aligned induced lane profiles and the sum of difference at the height of the Hsp90 peak was calculated (red area) (lower panel). Finally, the Hsp90 peak area was divided by the normalised and aligned EF-Tu peak area.

We determined the intensity profile of the uninduced (green) and induced lanes (orange) (**Figure 1B**, top panel) and normalised the induced lane profile to the uninduced lane profile using the EF-Tu peak of the uninduced lane as reference (100 %). Subsequently, we aligned the induced lane profile (green) to the uninduced lane profile (orange) using the first two left peaks (stars) of the uninduced lane profile as reference points (**Figure 1B**, middle panel). We deducted the uninduced lane profile (orange) from the normalised and aligned induced lane profile (green) and calculated the sum of difference at the height of the Hsp90 peak (red area) (**Figure 1B**, lower panel). Finally, we divided the Hsp90 peak area by the area of the normalised and aligned EF-Tu peak.

### The yield of recombinant human Hsp90β protein in E. coli using LB medium was insufficient for preparative purposes

We tested Hsp90 overproduction in LB medium varying the length of induction, the temperature, the OD_600_ at induction and the IPTG concentration to measure if any of these parameters modulated the levels of the chaperone.

After one and three days of induction no Hsp90 band appeared in the induced samples, none of the conditions led to sufficient Hsp90 overproduction (**Figure 2A and B**, lane 3-14). When induced for five days an Hsp90 band was apparent in several conditions (**Figure 2C**, lane 4, 5, 11 and 12). However, the yield was still insufficient for preparative purposes and the production levels were variable.

**Figure 2:**
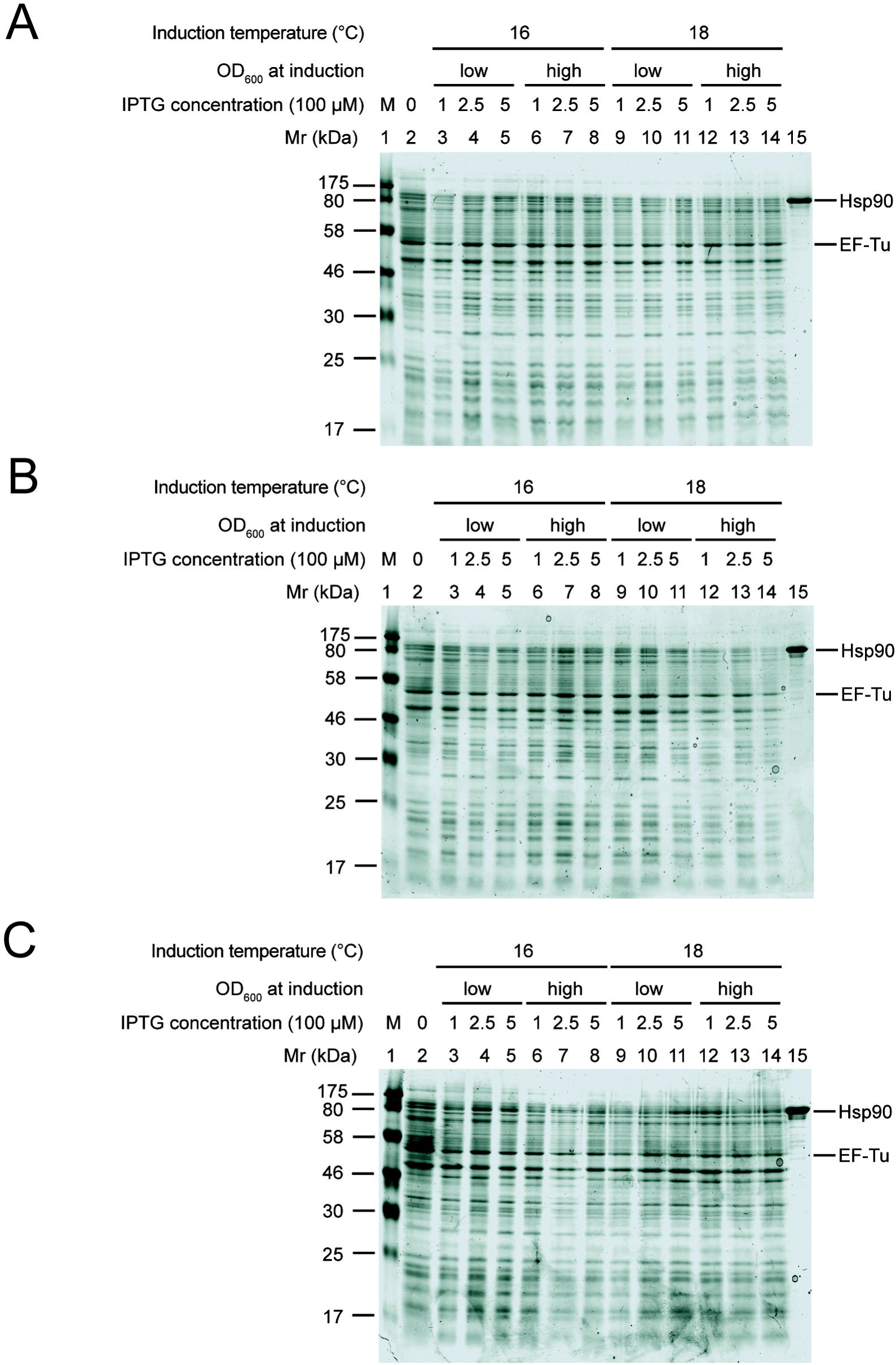
Hsp90 overproduction in LB medium lead to insufficient protein yield. Hsp90 overproduction after 1 day (A), 3 days (B), 5 days (C) of induction. Lane 1: protein marker, lane 2: uninduced sample, lane 3-8: protein production at 16°C, lane 9-14: protein production at 18°C, lane 3-5 and 9-11: induction at low OD_600_, lane 6-8 and 12-14: induction at high OD_600_, lane 3, 6, 9, 12: induction with 0.1 mM IPTG, lane 4, 7, 10, 13: induction with 0.25 mM IPTG, lane 5, 8, 11, 14: induction with 0.5 mM IPTG, lane 15: purified Hsp90 sample. EF-Tu intrinsic E. coli protein band was used for quantification of Hsp90 overproduction.

### 2x YT medium improved the yield of Hsp90 production

Since protein production in LB resulted in poor yields we hypothesised that the nutrient value of the medium may be a critical parameter in case of Hsp90. Therefore, we repeated the overproduction experiment in the richer 2x YT medium. We varied the length of induction, the temperature, the OD_600_ at induction and the IPTG concentration to determine if any these parameters affect Hsp90 production in 2x YT.

After one and three days of induction a new band appeared at the height of Hsp90 in the induced samples, suggesting that we could produce the chaperone (**Figure 3A and B**). The newly appearing Hsp90 band is stronger after three days of induction compared to one. Hsp90 production in 2x YT resulted in higher yields than in LB within the interval of the parameters we varied.

**Figure 3:**
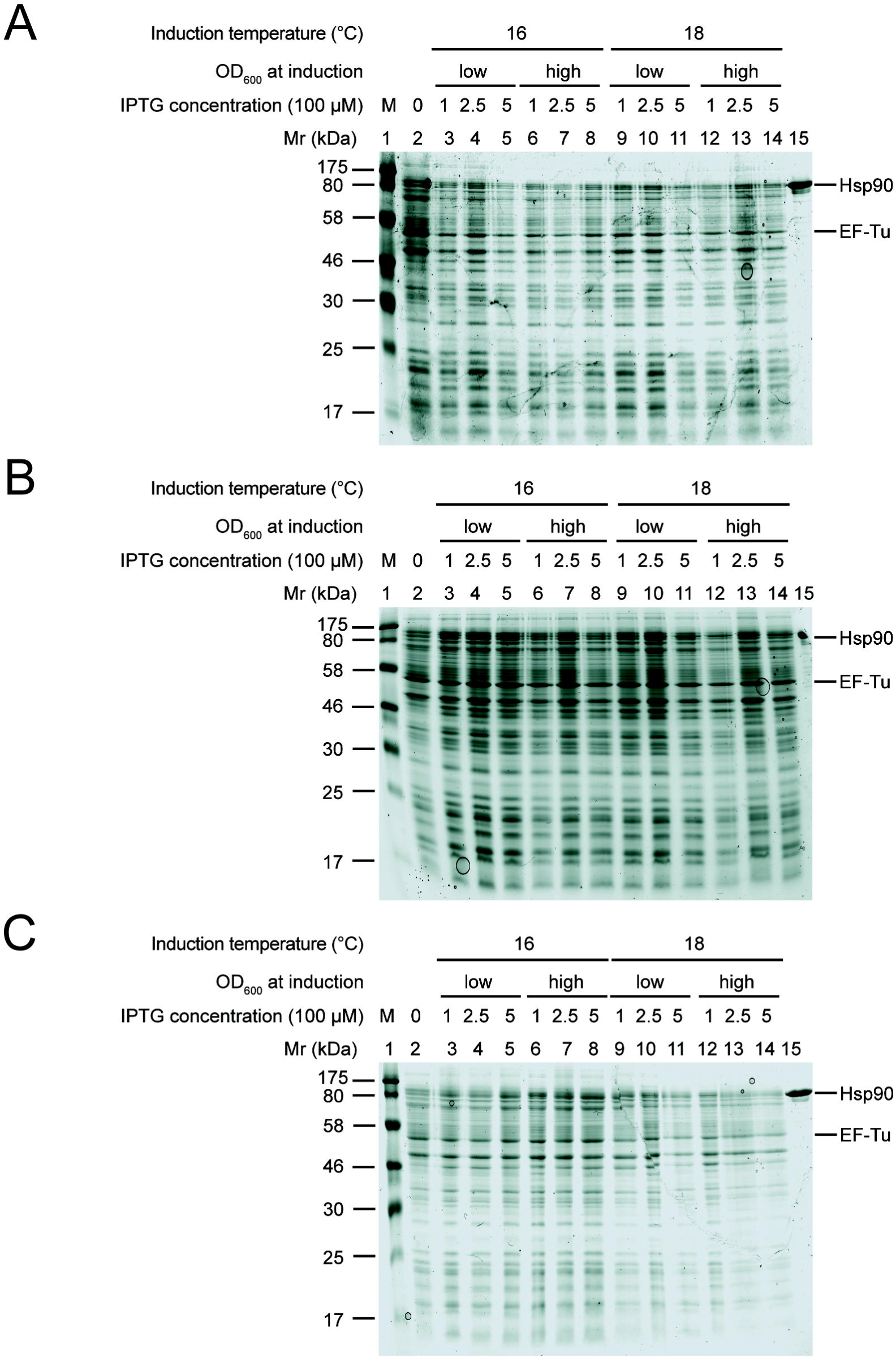
Hsp90 overproduction was improved but still insufficient in 2x YT compared to LB. Hsp90 overproduction after 1 day (A), 3 days (B), 5 days (C) of induction. Lane 1: protein marker, lane 2: uninduced sample, lane 3-8: protein production at 16°C, lane 9-14: protein production at 18°C, lane 3-5 and 9-11: induction at low OD_600_, lane 6-8 and 12-14: induction at high OD_600_, lane 3, 6, 9, 12: induction with 0.1 mM IPTG, lane 4, 7, 10, 13: induction with 0.25 mM IPTG, lane 5, 8, 11, 14: induction with 0.5 mM IPTG, lane 15: purified Hsp90 sample. EF-Tu intrinsic E. coli protein band was used for quantification of Hsp90 overproduction.

When induced for five days, we observed significant overproduction of the chaperone (**Figure 3C**) in certain cases (lane 7-9), whereas other conditions led to similar results to those observed on previous gels (lane 10, 12). Hsp90 production levels were variable in the different conditions. We concluded that Hsp90 overproduction resulted in higher yields in 2x YT after five days of induction compared to shorter induction times and culturing in LB medium, but the yields were still insufficient for preparative purposes.

### Recombinant human Hsp90β production had the best yield in the E. coli cells in TB medium

Since the richer 2x YT medium had a positive effect on Hsp90 levels we decided to test the protein production in TB medium which is higher in nutrients than 2x YT. We varied the same parameters as described previously.

After one day of induction, a strong band appeared at the height of Hsp90 in several conditions (**Figure 4A**, lane 3 and 11) indicating that overproduction in TB was notable. The conditions that resulted in high protein amount, however, were variable. Already after three days of induction in TB medium resulted all the conditions tested in a strong Hsp90 band (**Figure 4B**). We observed similar Hsp90 levels with five days of induction (**Figure 4C**). Thus, TB medium resulted in higher yields compared to both LB and 2x YT media, and after a shorter period of time.

**Figure 4:**
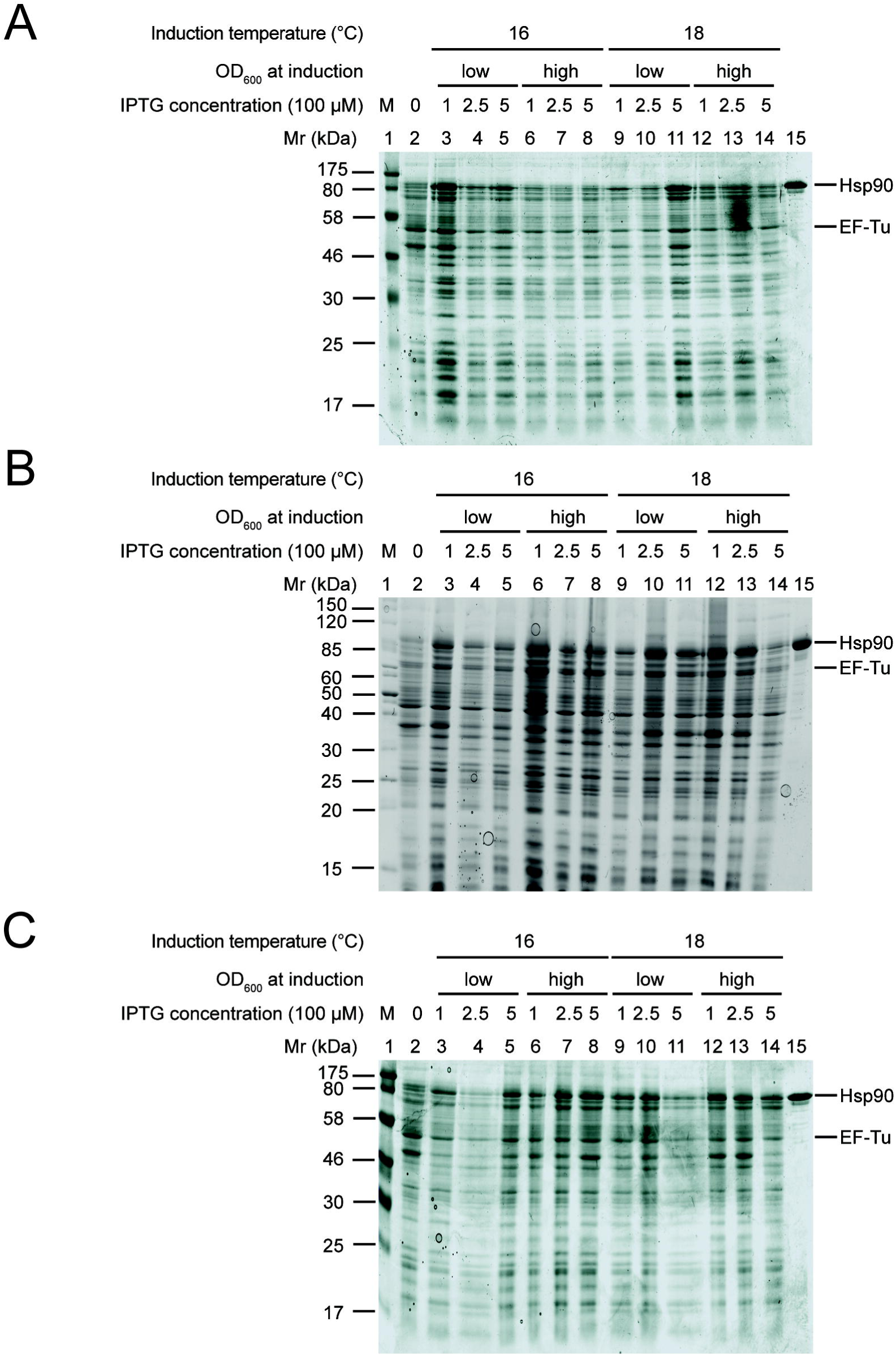
Hsp90 overproduction lead to the highest yields in TB medium. Hsp90 overproduction after 1 day (A), 3 days (B), 5 days (C) of induction. Lane 1: protein marker, lane 2: uninduced sample, lane 3-8: protein production at 16°C, lane 9-14: protein production at 18°C, lane 3-5 and 9-11: induction at low OD_600_, lane 6-8 and 12-14: induction at high OD_600_, lane 3, 6, 9, 12: induction with 0.1 mM IPTG, lane 4, 7, 10, 13: induction with 0.25 mM IPTG, lane 5, 8, 11, 14: induction with 0.5 mM IPTG, lane 15: purified Hsp90 sample. EF-Tu intrinsic E. coli protein band was used for quantification of Hsp90 overproduction.

### Quantification of overproduction gels revealed that the richness of the medium and the length of induction are the most significant parameters for Hsp90

To compare the protein production results of the different gels we estimated Hsp90 overproduction by gel densitometry as described in Figure 1B. In LB medium we observed low level of Hsp90 overproduction in each condition (**Figure 5A**). The tendency improved with the length of induction but even after five days the yields were insufficient for preparative purposes in all conditions.

**Figure 5:**
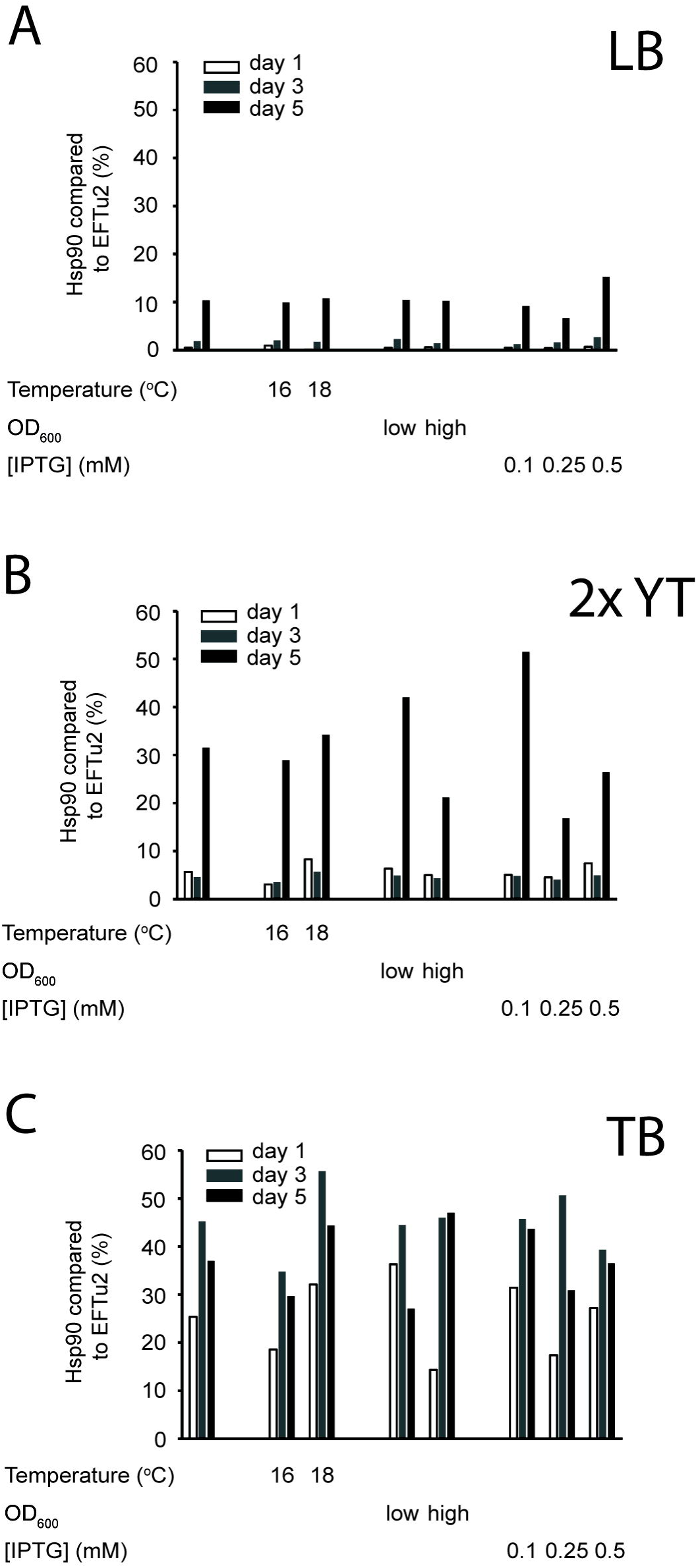
Quantification of Hsp90 overproduction in different media. (A)Summary of the quantification of Hsp90 overproduction in LB, after 1 day (white), 3 days (grey) and 5 days (black) of induction. Average of results of all conditions (1^st^ column group), average of results on 16°C and 18°C (2^nd^ and 3^rd^ column groups), average of results at low or high OD_600_ at induction (4^th^ and 5^th^ column groups), average of results at 0.1, 0.25 and 0.5 mM IPTG concentration (6^th^ to 8^th^ column groups). The length of induction is the only variable that influences Hsp90 production. (B) Summary of the quantification of Hsp90 overproduction in 2x YT. The length of induction is the only variable that influences Hsp90 production. The yield is higher in 2x YT compared to LB. The structure of the chart is as in (A). (C) Summary of the quantification of Hsp90 overproduction in TB. The length of induction is the only variable that has a significant effect on Hsp90 production. The yield is higher in TB compared to LB and 2x YT. The structure of the chart is as in (A).

We noted similar trends in case of 2x YT medium, the level of protein overproduction was insufficient and had similar yields after one and three days of induction (**Figure 5B**). After five days of induction in 2x YT, however, Hsp90 overproduction was still insignificant. Just like in LB, the other varied conditions did not modulate Hsp90 levels in this medium. In 2x YT medium the yield was higher in every condition compared to LB.

Hsp90 overproduction in TB was significantly higher than in the other two media. The length of induction had a significant effect on the Hsp90 yields, and here we reached good yields already after three days. The other tested parameters did not influence the Hsp90 yield in TB.

Overall we concluded that the quantification of the gels in Figure 2, 3 and 4 confirmed our observations about the overproduction gels (**Figure 5A, B and C**, respectively). We found that of the tested parameters, richness of medium used for culturing *E. coli* cells and the length of induction were the only two that had a significant effect on Hsp90 overproduction and should be considered in the future. Temperature, OD_600_ at induction and IPTG concentration used for induction did not significantly modulate the yields of the chaperone within the tested intervals.

### Recombinant human Hsp90β purified in one day

After successful protein overproduction, we set out to optimise a fast, efficient and trustworthy purification protocol to ensure excellent sample quality. This was necessary because like other multi-domain proteins, human Hsp90β is also prone to the degradation. Its accessible, flexible regions, especially the charged linker between the N-terminal domain and the middle domain is often targeted by proteases, causing N-terminal degradation. Widely used purification protocols include time-consuming purification steps (such as dialysis, size exclusion chromatography or both) that slow down the procedure which may have detrimental effects on protein quality (16-19).

To avoid degradation that might occur as a consequence of long purification procedures and eventual freeze-thaw steps, we set out to develop a condensed protocol for purification of Hsp90 in one day that results in highly pure and stable sample. We purified N-terminally His_6_-tagged Hsp90 protein by Ni-affinity chromatography, anion exchange and heparin affinity chromatography (**Figure 6A**). To ensure sample quality and avoid degradation we carried out the experiment in the presence of protease inhibitors at low temperature (0-4°C). To check for protein quality throughout the purification process we ran samples of the peak fractions of each column on sodium dodecyl sulfate polyacrylamide gel electrophoresis (SDS-PAGE) (**Figure 6B**). We observed a gradual gain in purity and loss in degradation products and impurities. After the last step the new Hsp90 purification protocol resulted in a protein sample free of any significant contaminations detectable by Coomassie staining. The combination of the three columns was necessary for high sample purity and reproducibility.

**Figure 6:**
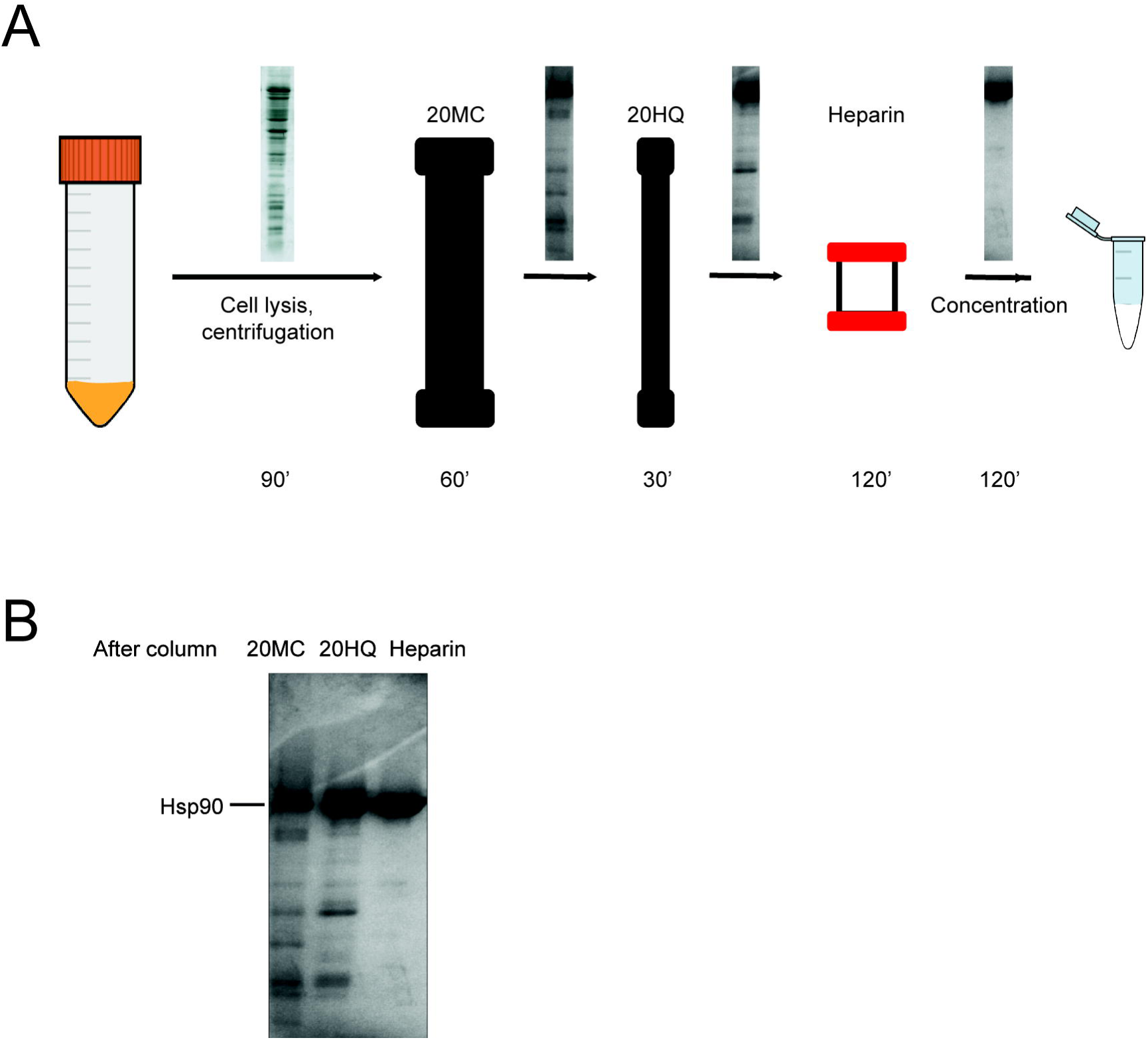
The Hsp90 sample was free of degradation products and impurities after the three-step purification procedure. (A) Schematic overview of the Hsp90 purification procedure. After cell disruption and centrifugation, the lysate was loaded first on a POROS 20MC Ni-column followed by a POROS 20HQ anion exchange column. Next, the sample was loaded on a heparin affinity chromatography column. Finally, the eluted protein sample was concentrated, buffer-exchanged and flash frozen. The time frame of the experiment is indicated on the bottom of the figure. The SDS-PAGE panels on the arrows show the purity of the samples after each purification step. (B) SDS-PAGE shows the purity of the Hsp90 sample after each purification step. Lane 1: peak fraction sample after POROS 20MC affinity chromatography purification step, lane 2: peak fraction sample after POROS 20HQ anion exchange purification step, lane 3: peak fraction sample after heparin affinity chromatography purification step.

Certain biochemical and biophysical experiments are carried out at a higher, physiological temperature (37°C) and for elevated time intervals (days or months). In case of proteins that are sensitive to proteolysis, incubation at high temperature for longer time can be detrimental for sample quality. To test the stability of the Hsp90 sample purified by the new protocol we incubated the protein at 4, 21 and 37°C and ran samples taken at 3, 18 and 24 hours on SDS-PAGE. We visualised Hsp90 in the gels first with Coomassie (**Figure 7A**) and subsequently, with more sensitive silver staining (**Figure 7B**).

**Figure 7:**
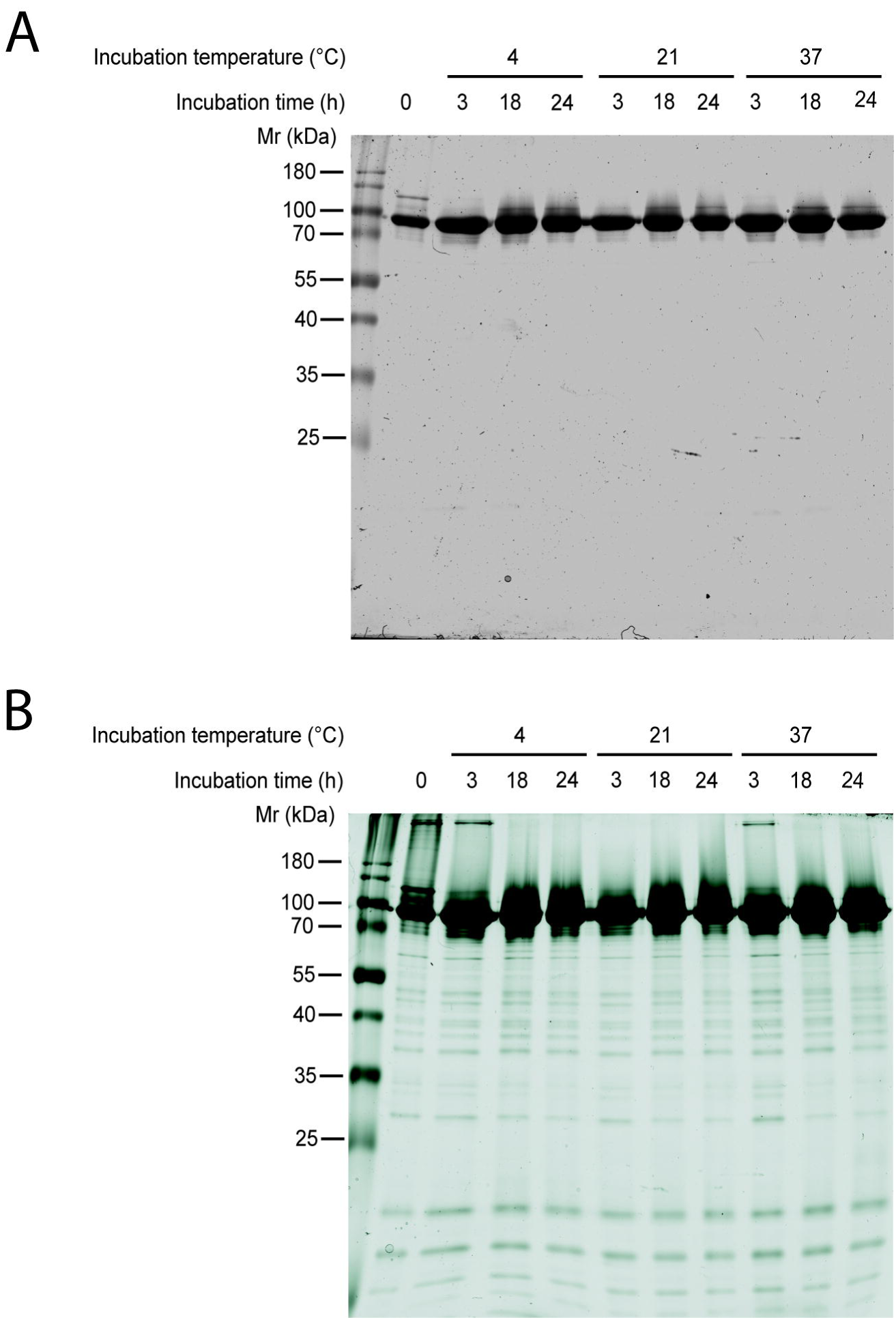
Purified Hsp90 sample incubated on physiological temperature for one day remained free of degradation products. (A) SDS-PAGE about the stability of the purified Hsp90 sample.Protein marker (lane 1), Hsp90 incubated for 0 hours (lane 2), Hsp90 incubated at 4oC (lane 3-5), 21oC (lane 6-8), 37°C (lane 9-11), Hsp90 incubated for 3 hours (lane 3, 6, 9), 18 hours (lane 4, 7, 10), 24 hours (lane 5, 8, 11). (B) Silver staining about the stability of the purified Hsp90 sample. Protein marker (lane 1), Hsp90 incubated for 0 hours (lane 2), Hsp90 incubated at 4°C (lane 3-5), 21oC (lane 6-8), 37°C (lane 9-11), Hsp90 incubated for 3 hours (lane 3, 6, 9), 18 hours (lane 4, 7, 10), 24 hours (lane 5, 8, 11).

No additional degradation products appeared in the incubated samples compared to the starting sample within the time frame of the experiment on the Coomassie-stained gels (**Figure 7A**). Therefore we concluded that Hsp90 purified using the new purification protocol was stable if incubated at 37°C up to 24 hours. Since Coomassie staining has limited sensitivity especially in the range of low molecular weight, we further examined silver staining to further examine the stability of the sample by silver staining (**Figure 7B**). Here, we observed impurities and/or degradation products in every sample but we did not see a systematic increase of any bands upon incubation at 37°C up to 24 hours.

This newly optimised, reproducible purification protocol enabled us to prepare a highly pure, stable and homogeneous sample within one day.

### Hsp90 purified by our method was properly folded, had the correct molecular weight and sedimentation coefficient

To reveal the folding status of proteins, we to analysed our Hsp90 samples with circular dichroism spectroscopy (CD) that can potentially reveal the secondary structure composition and folding status of the chaperone. The CD spectrum of Hsp90 revealed that the chaperone is mainly composed of α-helices (**Figure 8A**). We observed two minima at ∼209 nm and at ∼ 222 nm and a maximum at ∼ 194 nm in the spectrum which is in agreement with the previous findings (Prodromou et al, 1999).

**Figure 8:**
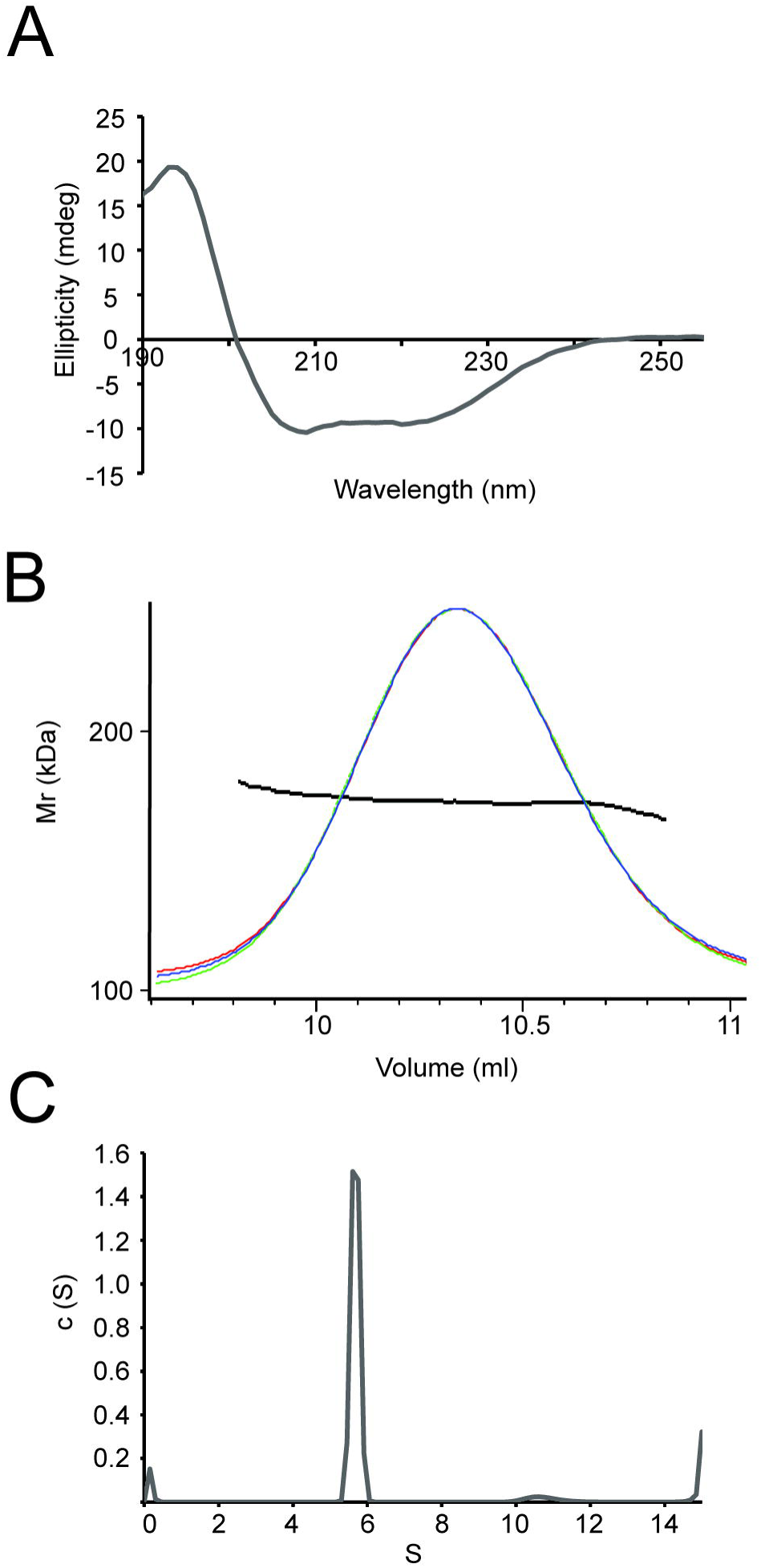
Purified Hsp90 sample folded, sedimented correctly and its molecular mass was appropriate. (A) CD spectrum of Hsp90 at 1.2 μM monomer concentration. The overall structure of the protein typically consists of α-helices. (B) The AUC spectrum of Hsp90 at 7.2 μM monomer concentration. Hsp90 dimers sediment with 5.6 S, whereas Hsp90 tetramers appear at 10.6 S. (C) SEC-MALLS spectra of the purified Hsp90 sample at 2.2 μM monomer concentration. The protein has the molecular weight of the dimer Hsp90 is 167.5 ± 13.3 kDa. The peak maximum is at 10.07 ml. Red line: light scattering, blue line: refractive index, green line: UV absorption at 220 nm, black dotted line: fit for molecular mass.

To ensure that our Hsp90 sample had the correct molecular weight and did not degrade during the purification process we measured its molecular weight using size exclusion chromatography - multi angle laser light scattering (SEC-MALLS) (**Figure 8B**). We observed a homogeneous peak at 10.07 ml that was fitted to 167.5 ± 13.3 kDa. This result corroborates with the expected size of two Hsp90 molecules (168.2 kDa), confirming that our sample is in a dimeric state at the concentration used in the experiment and also in agreement with data published earlier (Lepvrier et al, 2015, Moullintraffort et al, 2010). Taken together with the outcome of the stability experiments SEC-MALLS results suggested that our sample contained the intact full length Hsp90 dimer without degradation.

We measured the sedimentation coefficient of our Hsp90 sample to analyse sedimentation properties of the Hsp90 dimer. In the sedimentation profile of the chaperone we observed a peak at 5.6 S that corresponded to the dimer and a smaller peak at 10.6 S that was probably the tetramer molecule (dimer of dimers) (**Figure 8C**). The increasing fraction above 14 S suggested that the sample aggregated to some extent (but this is a typical phenomenon in case of analytical ultrancentrifugation (AUC) samples). Below 1 S we observed a small peak, that could have originated from impurities or degradation products.

## Discussion

Human Hsp90β is often produced in baculovirus system in insect Sf9 cells that is expensive, requires long preparation steps and is often difficult to scale-up (20-22). Our optimised recombinant human Hsp90β production protocol in E. coli provides an attractive alternative for the baculovirus system.

First, we systematically tested the production of recombinant human Hsp90β in *E. coli* cells to identify parameters that modulate the yield of the chaperone. We showed that the length of induction and the nutrient level of the medium had significant effect on Hsp90 overproduction, whereas temperature, IPTG concentration and OD_600_ at induction (within the tested intervals) did not significantly modify the yields of Hsp90. Induction for three days in TB medium resulted in good yield of the chaperone.

Moreover, we purified Hsp90 using a protocol that allowed us to finish within one day and resulted in well-folded, stable and pure sample. Since Hsp90 is extremely sensitive to proteolysis because of its flexible linkers, fast preparation increases sample quality, in addition to being cheaper and time-efficient. Throughout the purification procedure impurities and degradation products disappeared from the sample and final product resisted further degradation upon incubation on elevated temperature for long time intervals.

Previous Hsp90 purification protocols often contained time consuming preparation steps such as dialysis, size exclusion chromatography or both that may affect protein quality (16-19). Using the combination of Ni-affinity chromatography, anion exchange and the Hsp90-specific heparin column allowed us to get rid of degradation products and impurities in only three purification steps. The resulting sample was sufficiently pure and resistant to proteolytic degradation upon incubation on physiological temperature for up to one day, and showed expected biophysical properties (CD, SEC-MALLS).

In our AUC experiments human Hsp90 sedimented predominantly as a dimer with S=5.6 and a small fraction as a tetramer with S=10.6. It is notable that human Hsp90 is known to be in extended conformation (D_max_=26 nm) in the absence of nucleotide and co-chaperone p23 (10). Similarly to our results, earlier findings showed that in the absence of nucleotides, consequently in its extended conformation, yeast Hsp90 also had the sedimentation coefficient of 5.6 (23). These results indicate that these two proteins sediment similarly when in extended conformation. However, in the presence of ATP that stabilises compact conformation, yeast Hsp90 sedimented with the coefficient of 6.8, suggesting that structural rearrangements altered its sedimentation properties (23). In similar experiments Hsp90 sedimented with the coefficient of 6.1 (24). The variations between these results may be explained by the differences between the origin of the samples (porcine brain Hsp90 that has different post-translational modification pattern and was α mixture of a and β isoform vs Hsp90), moreover, by buffer, salt and reducing agent conditions.

In summary, we optimised Hsp90 overproduction and a fast and efficient purification protocol for the protein. The resulting sample was pure, properly folded and resistant to degradation, consequently it is suitable for biochemical and biophysical experiments.

## Acknowledgements

We are grateful to Ineke Braakman for continuous support. We thank Jonas M. Dörr for his help with CD experiments and data analysis. We thank Camilla de Nardis, Nadia O. Leloup and Remco N. P. Rodenburg, Deniz Ugurlar and Revina C. van Scherpenzeel for their help in setting up SEC-MALLS experiments and their support in data interpretation. We are grateful to Priyanka Sahasrabudhe for her comments on the manuscript.

## Author contributions

### Contribution to manuscript

The concept of the study originates from M. Radli and S.G.D. Rüdiger. M. Radli drafted the first version of the manuscript and prepared all the figures in the chapter. M. Radli and S.G.D. Rüdiger wrote the manuscript.

### Contribution to experiments

M. Radli tested Hsp90 production, ran the SDS-PAGE gels and quantified Hsp90 overproduction.

M. Radli optimised the purification of Hsp90 and purified all the protein used for the experiments in this chapter.

M. Radli tested the stability of Hsp90 on different temperatures.

M. Radli performed and analysed the analytical ultracentrifugation (AUC), SEC-MALLS and circular dichroism (CD) experiments.

D.B. Veprintsev supervised the AUC experiments, helped analyzing the data and interpreting the results.

S.G.D. Rüdiger supervised the experiments.

